# A practical phenotyping framework for root system architecture reveals enhanced root vigor in an *Aegilops tauschii*-derived wheat line

**DOI:** 10.64898/2026.01.28.702258

**Authors:** Sultan Md Monwarul Islam, Izzat Sidahmed Ali Tahir, Kinya Akashi

**Affiliations:** United Graduate School of Agricultural Sciences, Tottori University, 4-101 Koyama-Cho-Minami, Tottori City, Tottori Prefecture, 680-0945, Japan; Wheat Research Program, Agricultural Research Corporation, Wad Medani P.O. Box 126, Sudan; International Platform for Dryland Research and Education, Tottori University, 1390 Hamasaka, Tottori City, Tottori Prefecture, 680-0001, Japan; Faculty of Agriculture, Tottori University, 4-101 Koama-Chou-Minami, Tottori City, Tottori Prefecture, 680-0945, Japan

**Keywords:** wheat, Multiple Synthetic Derivative, MSD417, high temperature stress, arid region, seminal root angle, coleorhiza

## Abstract

The Multiple Synthetic Derivatives (MSD) population is a unique hexaploid wheat resource that captures extensive genetic diversity from *Aegilops tauschii* and exhibits wide variation in agronomic traits. However, root system architecture (RSA), a key determinant of resource acquisition and stress adaptation, remains poorly characterized in this population. Here, we established a practical phenotyping framework for RSA analysis and evaluated MSD417 as a representative genotype. A two-dimensional cultivation platform enabling continuous imaging of seedling root growth under controlled conditions was established to quantify RSA traits and their responses to high temperatures. MSD417 was compared with its recurrent parent, Norin 61 (N61). Under controlled conditions, MSD417 displayed greater total root length, root system width, and convex hull area than N61, indicating enhanced early root vigor. This genotype also exhibited a wider seminal root angle, suggesting improved horizontal soil exploration while maintaining root depth. High-temperature treatment reduced overall root growth and minimized genotypic differences, indicating that temperature stress constrains RSA expression. Microscopic observations further revealed a lower height-to-width ratio of coleorhiza tissue of MSD417, suggesting restricted downward expansion. Collectively, this study establishes a practical framework for RSA phenotyping and demonstrates the potential of *Aegilops tauschii*-derived germplasm to enhance wheat root-related adaptive traits.

## Introduction

Wheat (*Triticum aestivum* L.) holds major agricultural importance as a staple food for a large portion of the global population, accounting for 18% of total dietary calories globally **(Erenstein et al. 2022)**. Climate change is expected to threaten wheat production, resulting in an estimated 16% reduction over the next three decades, particularly in the tropical regions of Africa and Southern Asia, where wheat is consumed daily as a staple food **(Pequeno et al. 2021)**. Wheat is highly sensitive to temperatures above the optimum range, and global wheat yield could drop by 6% for each degree-Celsius temperature rise **(Asseng et al. 2015; Farhad et al. 2023)**.

Root system architecture (RSA) refers to the set of traits (e.g., root length, number, angle, and distribution along the soil profile) used to characterize the structural features of the root system **(Lynch 2022; Cope et al. 2024)**. RSA is influenced by a combination of internal (e.g., underlying genetics and developmental influences) and external factors such as environmental causes (e.g., climate, physical and chemical properties of the soils, and biotic influence), leading to genotype-by-environment interactions that greatly affect root system architecture **(Lynch 2022; Kang et al. 2024)**. The plastic nature of the root system and its development allow plants to adapt to changing environments **(Lynch 2022)**, and it has been proposed as an important breeding target for a ‘second green revolution’ for the development of climate-resilient crops **(Lynch 2007; Ober et al. 2021)**.

Multiple Synthetic Derivatives (MSD) is a unique population of synthetic wheat that harbored a large genetic variation in *Aegilops tauschii* in a background of hexaploid wheat **(Tsujimoto et al. 2015; Elbashir et al. 2017; Gorafi et al. 2018, 2025)**. The MSD population was initially developed by crossing and backcrossing 43 different primary synthetic wheat lines with the Japanese wheat cultivar ‘Norin 61’ (N61). Successive self-pollination of the population up to BC_**1**_F_**5**_ generation resulted in the development of representative genotypes collectively called the MSD panel **(Gorafi et al. 2018, 2025)**. The genotypes in the MSD population show diverse morphology, agronomic traits, and their environmental responses, including heat, drought, and low soil nutrients such as nitrogen and phosphorus **(Tsujimoto et al. 2015; Elbashir et al. 2017; Gorafi et al. 2025)**. Based on the high field performance in Sudan, one of the hottest wheat-growing areas in the world, five selected MSDs genotypes (MSD296, MSD34, MSD392, MSD417, and MSD54) were further analyzed for their physiological and molecular responses to high temperature in a controlled environment, revealing that the diversity in the agronomic traits was associated with distinct molecular responses **(Matsunaga 2022)**. Among these genotypes, MSD417 exhibited one of the highest performances across various traits, including fertility, thousand-kernel weight, harvest index, and yield. These findings suggest that the MSD population is a useful genetic material for future breeding programs. However, features of the root structures and their environmental responses have not been examined to date.

Evaluating RSA traits is often challenging because access to the root is limited, typically requiring laborious, costly, and time-consuming procedures, which constrain phenotypic analysis. Nevertheless, fully exploiting variability in RSA traits could yield substantial benefits **(Johnson et al. 2001; Trachsel et al. 2011; Wasson et al. 2014; Maqbool et al. 2022; Cope et al. 2024)**. In-field measurements of the root system tend to be destructive **(Trachsel et al. 2011)**, thus mostly undertaken at a single time-point for an individual plant, and are often labor-intensive or capture only a portion of the RSA. RSA measurements in controlled conditions typically lead to more reproducible phenotypic results **(Poorter et al. 2016)**. Various analytical platforms have been used to assess RSA in soil-based research conducted in controlled environments, including rhizoboxes made of wood, glass plates, metal boxes, and transparent polyvinyl chloride tubes **(Schuurman and Goedewaagen 1971; Böhm 1979; Neumann et al. 2009)**. On the other hand, non-soil-based systems, such as transparent pots **(Richard et al. 2015)**, germination paper **(Gioia et al. 2016)**, agar plates **(Nagel et al. 2020)**, GLO-Roots **(Nagel et al. 2012)**, GrowScreen-Rhizo **(Rellán-Álvarez et al. 2015)**, and paper or cloth pouches **(Shorinola et al. 2019; Chen et al. 2020)** enable non-invasive time-course analyses of root development and easier root sampling. However, these systems often require investment in specialized devices. In the present study, we developed a simple and cost-effective two-dimensional platform for RSA analysis and used it to evaluate MSD417, a candidate model for heat-tolerant wheat genotypes in the MSD population.

## Materials and Methods

### Plant material

Seeds of bread wheat cultivar ‘Norin 61’ (hereafter referred to as N61), and MSD genotype MSD417 were kindly provided by Prof. Hiroyuki Tanaka (Faculty of Agriculture, Tottori University, Tottori, Japan) and Dr Yasir Serag Alnor Gorafi (Arid Land Research Center, Tottori University, Tottori, Japan), respectively. The seeds were multiplied in pots at a greenhouse facility in the Faculty of Agriculture, Tottori University, Tottori, Japan, and fully mature seeds were harvested in 2024 and used for the experiments.

### Construction of the root growth panel

Wheat plants were grown on a custom-made, two-dimensional root growth panel based on the protocols described by **(Shorinola et al. 2019; Chen et al. 2020)** with the following modification. First, two sheets of paper towel (Kimtowel, a four-layer pulp, unbleached, Nippon Paper Crecia, Tokyo, Japan) were cut into 34 cm in height and 30 cm in width by scissors, then the bottom corners of both sides were trimmed off to remove rectangles measuring 3 cm in height and 3 cm in width (Fig. 1a). The paper towels were soaked in a 2000-fold diluted HYPONeX nutrient solution (Hyponex Japan, Osaka, Japan) and then placed on the surface of an acrylic board (31 cm height, 30 cm width, and 0.5 cm thickness) to extend 3 cm below the bottom edge of the acrylic board. A 33 cm × 30 cm sheet of black calico cloth was laid on top of the paper towel. The cloth and paper towel were attached to the acrylic board by wrapping them with three 1 mm-wide No. 18 rubber bands (Daiso Industries, Higashi-Hiroshima, Hiroshima, Japan) horizontally, 2 cm below the top edge of the acrylic board (Fig. 1b). To prevent moisture on the surface of the calico cloth from evaporating, a high-density polyethylene sheet (L-501-2, 0.05 mm thickness, As One, Osaka, Japan) was cut into a 31 × 30 cm square and placed over the cloth. This allowed roots to grow between the polyethylene sheet and the calico cloth; the latter served as a supporting substance for root growth. To prevent light exposure to the roots, a second black calico cloth measuring 33 × 30 cm was placed over the polyethylene sheet, extending up to 2 cm above the top edge of the acrylic board. The 2 cm protruding part of the cloth was folded over the top of the acrylic board to prevent light from entering the stack, while the center of the folded cloth was cut out to create a 3 cm wide window for shoot emergence. Four 3 cm wide binding clips were placed at the corners of the board to secure the entire system (Fig. 1c). Hereafter, this system is referred to as the root growth panel.

**Fig. 1.**
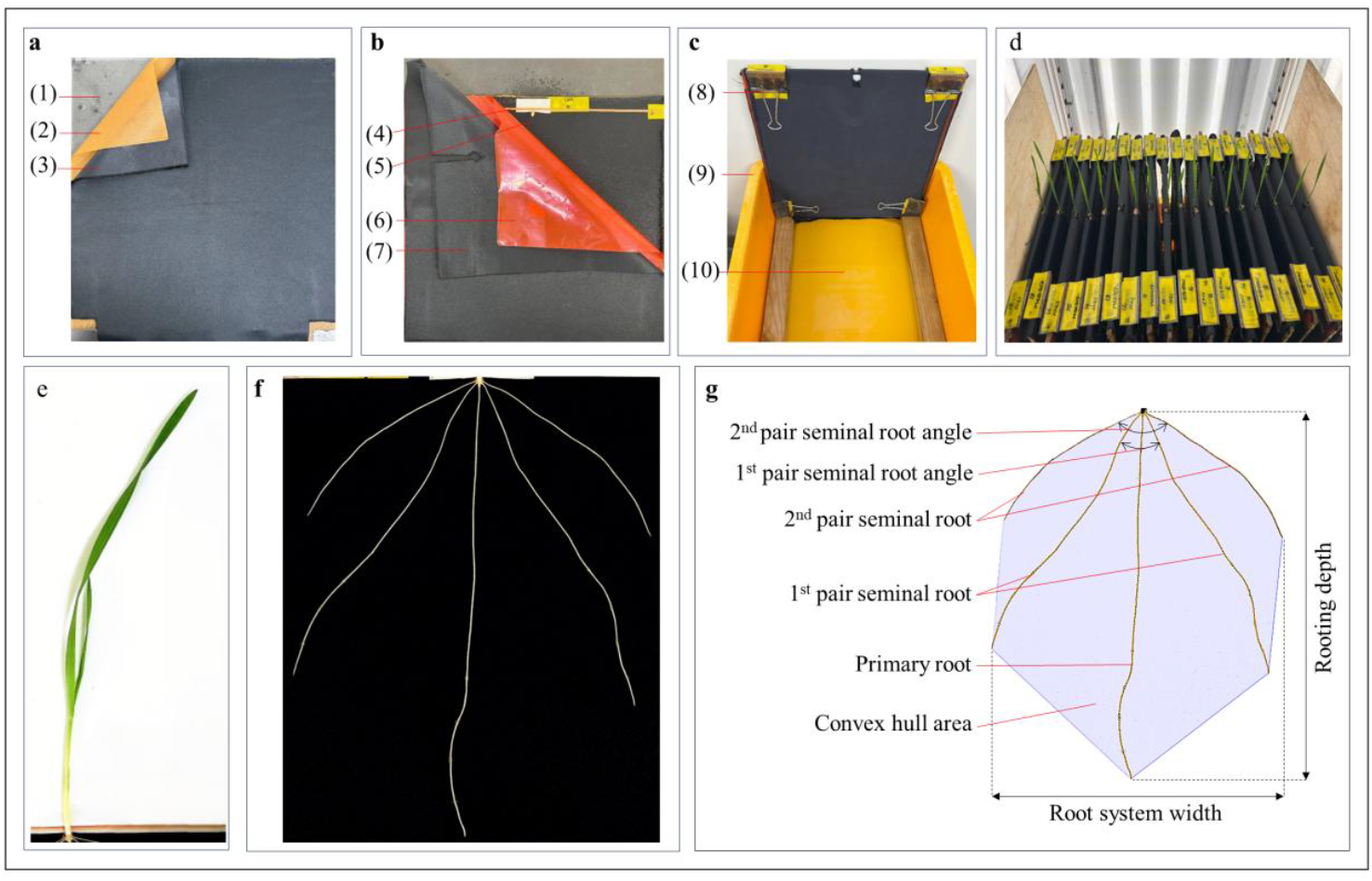
Two-dimensional root growth panel for analysing root system architecture. (a) A stack on the back side of the panel composed of an acrylic board (1), sheets of a kimtowel (2), and a black calico cloth (3). (b) A placement and further stacking onto the back side stack by rubber bands that secured the seed (4), a pre-germinated seed (5), a polyethylene sheet (6), and a second black calico cloth (7). (c) Installation into the container. The stacked panel was secured at four corners with clips (8) and placed in a plastic tray (9) that was filled with the nutrient-mixed water solution (10). (d) The vertically stacked panels in a growth chamber. (e) A photograph of a representative MSD417 shoot at day 8 under control conditions. (f) A representative image of root growth of N61 at day 8 under high-temperature conditions. (g) An image of root tracing by ImageJ plugins with SmartRoot and its root traits

To accommodate the root growth panels in a growth chamber, two pieces of wood (48 cm in length, 4 cm width, and 3 cm in height) were placed along the inner bottom of a plastic tray (50 cm in length, 35 cm in width, and 20 cm in height) (Fig. 1c), and filled with the 2000 folds-diluted HYPONeX solution to a depth of 3 cm. The 18 panels were stacked vertically on top of the woods in the tray (Fig. 1d), ensuring that the paper towels protruding downward from the panels were submerged in the solution. Finally, the whole unit, including the growth panels, was covered (except the upper middle part of the growth panels) with the same calico cloth (45 cm × 45 cm), followed by a high-density polyethylene sheet (45 cm × 45 cm) to protect water evaporation from the growth panels. The unit was placed in a growth chamber (14/10 h light/dark regime, 22 °C/18 °C day/night temperatures, light intensity of approximately 350 µmol m^−2^ s^−1^, and relative humidity of 50-60%).

### Plant growth conditions

The seeds were placed on top of two 85 mm-diameter filter papers (Filter paper type-2, Advantec, Tokyo, Japan) in a 90 mm-diameter petri dish and imbibed with 10 ml of tap water. Another piece of the same filter paper was used to cover the seeds. The petri dish was capped with a transparent lid, wrapped with aluminum foil, and incubated at 4 °C for two days. The petri dish was incubated at 25 °C in the dark for another day. The uniformly germinated 12 seeds of each genotype were placed at the central position of the rubber band in the root growth panels and covered by two pieces of filter paper (3 cm × 2 cm), which were moisturized in advance with a 2000-fold HYPONeX solution to prevent the seeds from drying (Fig. 1b). The panels were incubated under normal temperature conditions (14/10 h light/dark regimes, 22 °C/18 °C day/night temperatures). After four days, half of the panels were transferred to a high-temperature chamber with day/night temperatures of 42 °C/18 °C for an additional four days, while the control plants were maintained at normal temperature conditions (22 °C/18 °C).

### Root imaging and biomass measurement

To visualize the root system architecture, the polyethylene sheets and light-blocking black calico cloth were removed from the root growth panel, and the root systems and shoot morphology were photographed using a bench-top photograph system (model ET16 Plus, CZUR Tech, Shenzen, China). After taking root image, the 2000-fold HYPONeX solution was evenly sprayed onto the roots, followed by reassembling the supporting black calico cloth and the panels for further growth. This image acquisition was performed daily. On the eighth day, the root and shoot were harvested separately, weighed on an electric balance to record fresh weight, and then dried for three days in a drying oven at 80 °C to record their dry weight.

### Root image processing and statistical analysis

All root images were processed using ImageJ (version 1.54g) plugins with SmartRoot (Version 4.21) **(Lobet et al. 2011)**. Rooting depth, root system width, individual root length, seminal root angle, and shoot height were measured. Data were analyzed and visualized in the R statistical environment (version 4.5.0, R Core Team, 2025) using a set of custom-made R scripts deposited in Supplementary Document S1. Tukey’s method was applied for multiple comparisons.

## Results

### Root biomass allocation and root system architecture

The germinated wheat plantlets were grown in root growth panels under controlled conditions (22/18 °C day/night) for four days. Then, half of the plantlets were transferred to a high-temperature growth chamber with day/night temperatures of 42/18 °C for an additional four days to undergo high-temperature treatment. In N61, the primary roots reached approximately two-thirds of the growth panel depth in eight days, accompanied by typically four seminal roots, with a primary root extending diagonally downward (Fig. 2a). High temperature resulted in the growth inhibition for both the primary and the seminal roots in N61 (Fig. 2b). MSD417 showed a similar pattern for root growth, but a tendency toward longer roots and a broader root system width was observed (Figs. 2c, d).

**Fig. 2.**
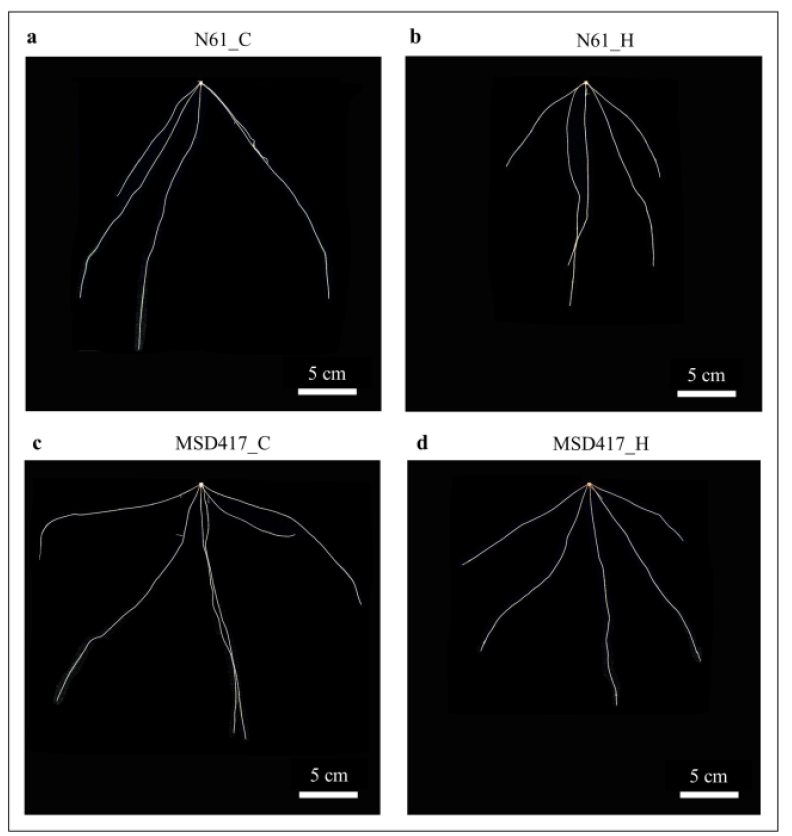
Representative images of N61 and MSD417 in contrasting temperature conditions at day eight. (a) N61_C-N61 in control condition, (b) N61_H-N61 in high-temperature condition, (c) MSD417_C-MSD417 in control condition, and (d) MSD417_H-MSD417 in high-temperature condition

To evaluate root system characteristics quantitatively, trait parameters were measured as defined in Table 1 and Fig. 1g. Root dry weight (RDW) differed significantly among genotypes and treatment conditions (Fig. 3a). Under control conditions, MSD417 exhibited significantly higher RDW than N61, both under control and high-temperature conditions, as well as its RDW under high-temperature conditions. In contrast, no statistical difference in shoot dry weight (SDW) was observed between genotypes or temperature conditions (Fig. 3b). Consequently, the root-to-shoot ratio (RSR) was the highest in the MSD417 under control conditions (mean value of 0.81) compared to the other genotype/condition combinations (mean values in the range of 0.62–0.67) (Fig. 3c), suggesting that the MSD417 genotype possesses a preferential allocation of biomass to the root system under non-stress conditions.

**Table 1.** Traits measured in the root growth panels.

| Traits | Abbreviation | Unit | Note |
| --- | --- | --- | --- |
| Rooting depth | RD | cm | Vertical length from root base to root tip |
| Shoot height | SH | cm | Length between the shoot base and tip |
| Root dry weight | RDW | mg |  |
| Shoot dry weight | SDW | mg |  |
| Root-to-shoot ratio | RSR |  | Root dry weight / shoot dry weight |
| Total root length | TRL | cm | Total length of all seminal roots and primary root |
| Specific root length | SRL | cm mg<sup>-1</sup> | Total root length/root dry weight |
| Primary root length | PRL | cm | Length of primary root |
| 1<sup>st</sup> pair seminal root length | FPSRL | cm | Mean length of the first two seminal roots |
| 2<sup>nd</sup> pair seminal root length | SPSRL | cm | Mean length of the third and fourth seminal roots |
| 1<sup>st</sup> pair seminal root angle | FPSRA | degree | Angle between the first two seminal roots |
| 2<sup>nd</sup> pair seminal root angle | SPSRA | degree | Angle between the third and fourth seminal roots |
| Root system width | RSW | cm | Maximum horizontal spread of the root system |
| Convex hull area | CHA | cm<sup>2</sup> | A convex polygon drawn to enclose all the outermost points of the roots. |

**Fig. 3.**
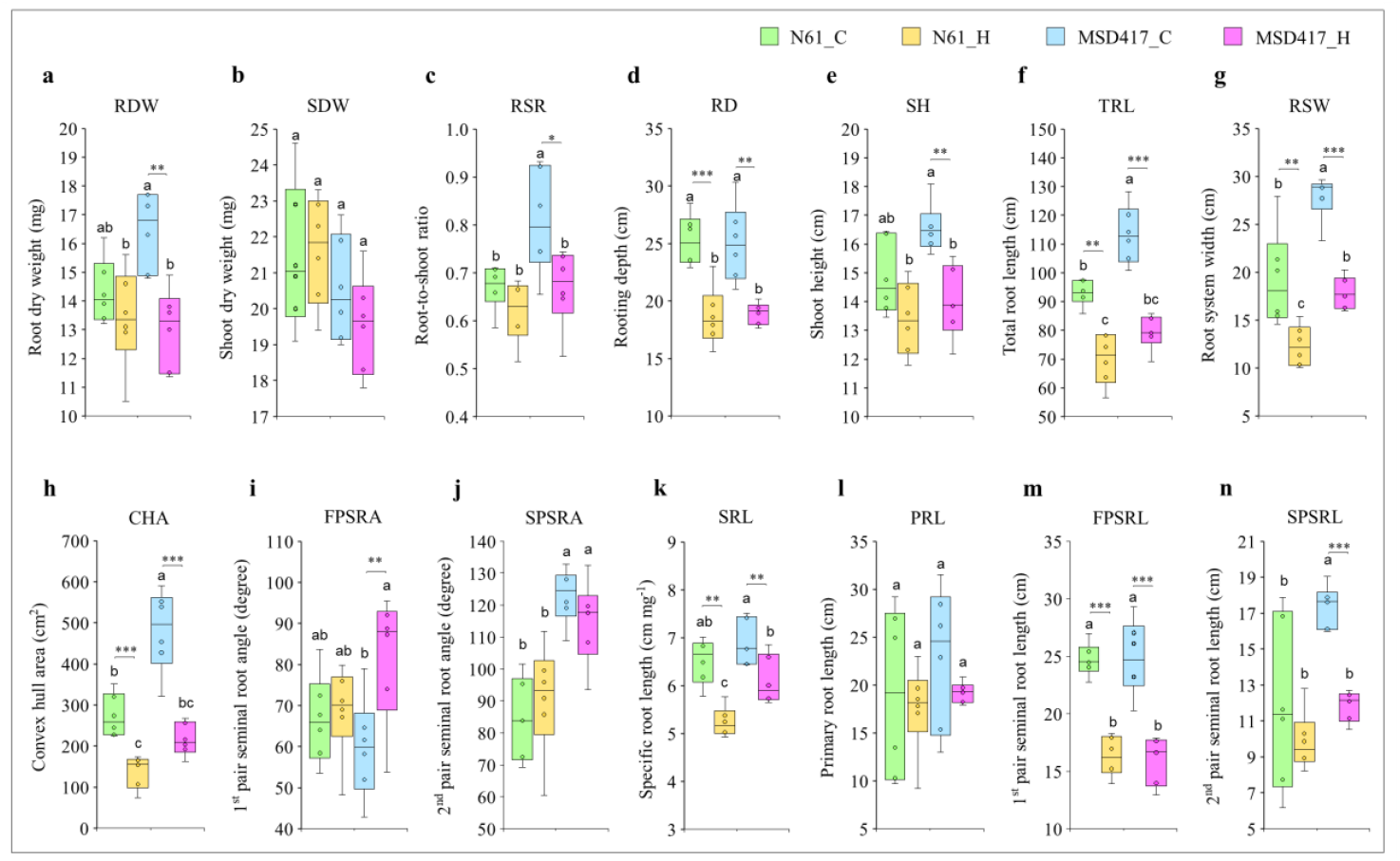
Variation in root and shoot morphological traits in contrasting temperatures of N61 and MSD417. The legend at the top of the panels shows four genotype-by-environment combinations: N61_C, N61_H, MSD417_C, and MSD417_H, representing N61 in the control condition, N61 in the high-temperature condition, MSD417 in the control condition, and MSD417 in the high-temperature condition, respectively. (a) root dry weight, (b) shoot dry weight, (c) root-to-shoot ratio, (d) rooting depth, (e) shoot height, (f) total root length, (g) root system width, (h) convex hull area, (i) first pair seminal root angle, (j) second pair seminal root angle, (k) specific root length, (l) primary root length, (m) first pair seminal root length, and (n) second pair seminal root length. Data for N61 and MSD417 were compared using Tukey’s HSD post hoc test; different letters indicate significant differences among genotypes under contrasting temperature conditions. Data for the same genotypes raised under contrasting temperature conditions were compared using Student’s t-test (p < 0.05, p < 0.01, p < 0.001; n = 6). Abbreviations of the trait parameters are shown in Table 1

Despite these differences in biomass allocation, traits such as rooting depth (RD) and shoot height (SH) did not differ significantly between the two genotypes after eight days (Figs. 3d, e). This indicated that the genotypic differences in root traits were not associated with vertical soil exploration. Under high-temperature conditions, both genotypes showed significant reductions in various traits. For instance, the RD was significantly affected by high temperature in both genotypes, resulting in 26% and 24% reductions in N61 and MSD417, respectively, compared with the control conditions (Fig. 3d). Since no genotypic differences in RD were observed, subsequent analyses focused on geometric parameters describing the spatial configuration and distribution of the root system.

### Geometric configuration of the root system

To further characterize root system architecture, geometric traits describing root length distribution and spatial expansion were analyzed. Under the control condition, total root length (TRL) was longer in MSD417 (113.25 ± 9.12 cm) than in N61 (93.02 ± 3.98 cm), while high temperature decreased the TRL in both genotypes (Fig. 3f). Root system width (RSW) and convex hull area (CHA) were also significantly larger in MSD417 (Figs. 3g, h), indicating enhanced horizontal expansion in MSD417. Notably, while the angle between the first pair of seminal roots did not differ significantly between the two genotypes (Fig. 3i), the angle between the second, outermost pair of seminal roots was larger in MSD417 (122.96 ± 7.85 and 114.91 ± 11.92 degree under control and high temperature conditions, respectively), in comparison to those of N61 (84.39 ± 11.46 and 90.72 ± 15.79 degree, respectively) (Fig. 3j), suggesting that the wider root system in MSD417 is at least partly attributable to the increased divergence of the outermost seminal roots.

### Contribution of the individual root components

Specific root length (SRL) showed no significant change under the control conditions of N61 and MSD417 (Fig. 3k); however, SRL decreased significantly under the high-temperature conditions for both genotypes. Primary root length (PRL) did not differ significantly among conditions or genotypes (Fig. 3l), and the length of the first pair seminal root (FPSRL) was significantly shorter under the high-temperature conditions for both genotypes (Fig. 3m). In contrast, the length of the second pair seminal root (SPSRL), corresponding to the outermost seminal roots, was significantly greater for the MSD417 under the control conditions (17.37 ± 1.06 cm) than for N61 (11.89 ± 4.30 and 9.84 ± 1.48 cm under control and high-temperature conditions, respectively), and MSD417 high-temperature conditions (11.85 ± 0.77 cm) (Fig. 3n), highlighting the uniqueness of this trait in MSD417 genotype.

### Temporal dynamics of seminal root development

Time course analysis further confirmed the architectural divergence between the two genotypes. The elongation of PRL and FPSRL did not differ significantly between the N61 and MSD417 genotypes under either control or high-temperature conditions (Figs. 4a, b). By contrast, SPSRL was consistently greater in the MSD417 under the control conditions throughout the examined period compared to the control of N61 and the high-temperature conditions of both genotypes (Fig. 4c), suggesting that early vigorous development of the outermost seminal roots during the seedling stage is a defining characteristic of the MSD417 genotype.

**Fig. 4.**
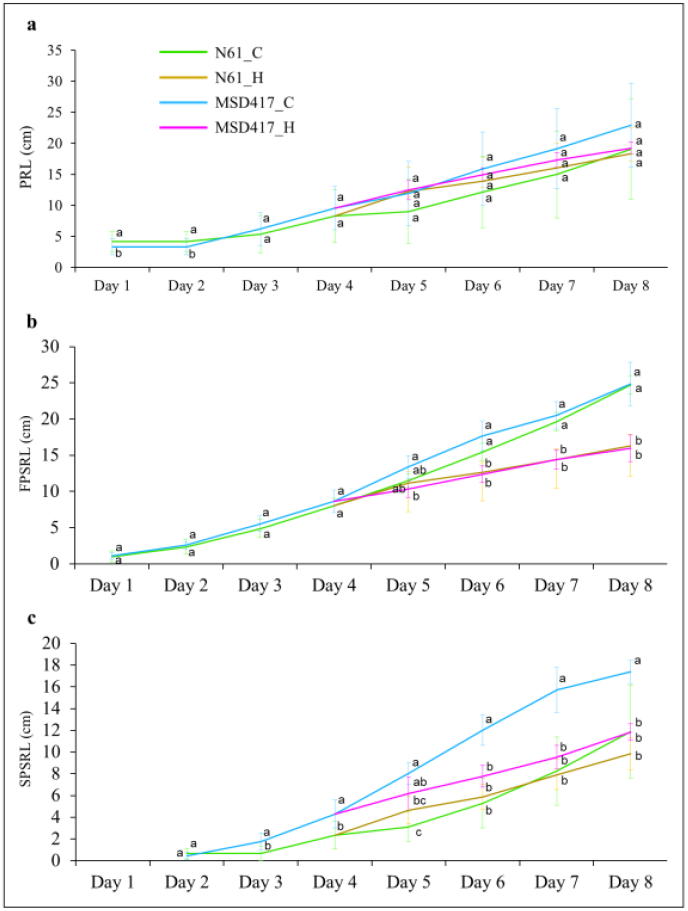
Line graphs showing the variation in the development of different root traits of N61 and MSD417 over different days after germination in contrasting temperatures. (a) primary root length, (b) 1^st^ pair seminal root length, and (c) 2^nd^ pair seminal root length. The legend at the top of the panels shows four genotype-by-environment combinations: N61_C, N61_H, MSD417_C, and MSD417_H, representing N61 in the control condition, N61 in the high-temperature condition, MSD417 in the control condition, and MSD417 in the high-temperature condition, respectively. PRL- primary root length, FPSRL- 1^st^ pair seminal root length, and SPSRL-2^nd^ pair seminal root length. Data for the N61 and MSD417 were compared using Tukey’s HSD post hoc test; different letters denote significant differences among genotypes under contrasting temperature conditions (n = 6)

### Multivariate structure of root system architecture

To visualize relationships among traits and to gain insight into the main sources of variation among genotypes under contrasting temperatures, principal component analysis (PCA) was performed (Fig. 5). The first two principal components (PC1 and PC2) collectively explained 73.6% of the total variance, with PC1 accounting for 52.6% and PC2 for 21.0%. The PCA scores largely separated the control and high-temperature conditions along PC1, with the control plants of both genotypes clustered on the positive side, whereas the high-temperature-stressed plants were positioned on the negative side. Separation of genotypes was also observed; plants in the MSD417 control condition clustered in the higher PC1 region, while those in N61 tended to cluster near the origin or scattered toward the PC2-negative plane. Under high-temperature conditions, MSD417 tended to cluster in the PC2-positive region, whereas N61 showed a tendency toward lower PC2 values.

**Fig. 5.**
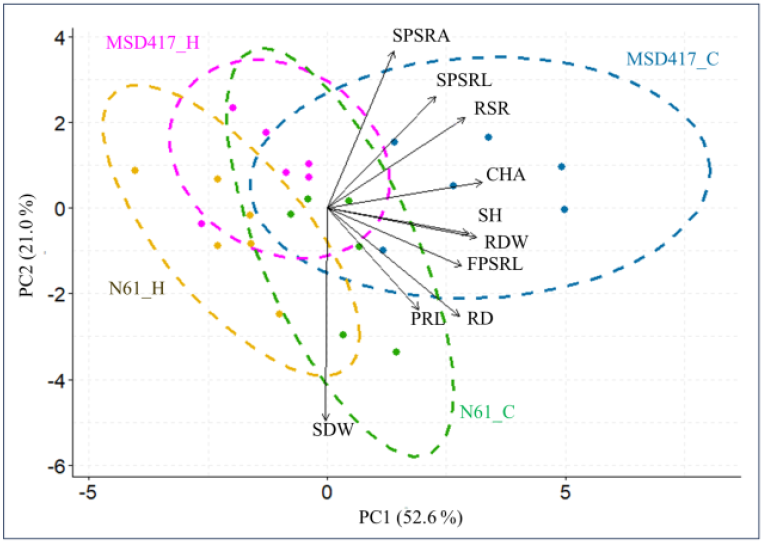
Principal component analysis (PCA) biplot of different traits of N61 and MSD417 in the contrasting temperature conditions. Colored dots represent the scores of plant individuals (n = 6) for N61 in the control condition (N61_C), N61 in the high-temperature condition (N61_H), MSD417 in the control condition (MSD417_C), and MSD417 in the high-temperature condition (MSD417_H), respectively. Vectors in black color indicate directions and strength of the loadings of the trait parameters as shown in Table 1.

The loading vectors showed that most traits had strong positive loadings on PC1, suggesting that these traits were strongly associated with the control condition and were inhibited under the high-temperature condition (Fig. 5). Interestingly, SPSRA and SPSRL had higher PC2 values than the other parameters, suggesting that the larger angle and length of the second pair seminal roots (SPSRL) are associated with MSD417 genotypes. It was also found that RSR and CHA were associated with the MSD417 under control conditions, suggesting a trend toward a higher root-to-shoot ratio and a larger root convex hull area in this genotype. By contrast, the loading of SDW showed a strongly negative PC2 value (Fig. 5), consistent with the trend of higher shoot dry weight in N61 compared to MSD417 (Fig. 3b).

### Root traits correlations of the two genotypes at different temperatures

To examine relationships among root parameters, correlations were calculated for each genotype-temperature condition combination (Fig. 6). In the N61 genotype under the control condition, many trait parameter combinations showed positive correlations, including RD with RDW, SDW, and RSW; SH with SDW, PRL, and RSW (Fig. 6a). However, SPSRL, SRL, RSR, and RSW showed negative correlations with several other root traits, including SH and SDW. On the other hand, under the high-temperature condition N61, while the majority of trait pairs correlated positively, SPSRA remained inversely correlated with most traits, and SRL showed negative correlations with RSR and RD (Fig. 6b). In the MSD417 genotype, distinct trends in parameter correlations were observed compared with N61. Under the control condition, SDW showed negative correlations with RDW, RSR, SH, and SPSRL (Fig. 6c). In contrast, under the high-temperature condition, new sets of strong negative correlations were observed, including SRL with RDW, RSR, and SH, as well as SDW with SPSRA (Fig. 6d).

**Fig. 6.**
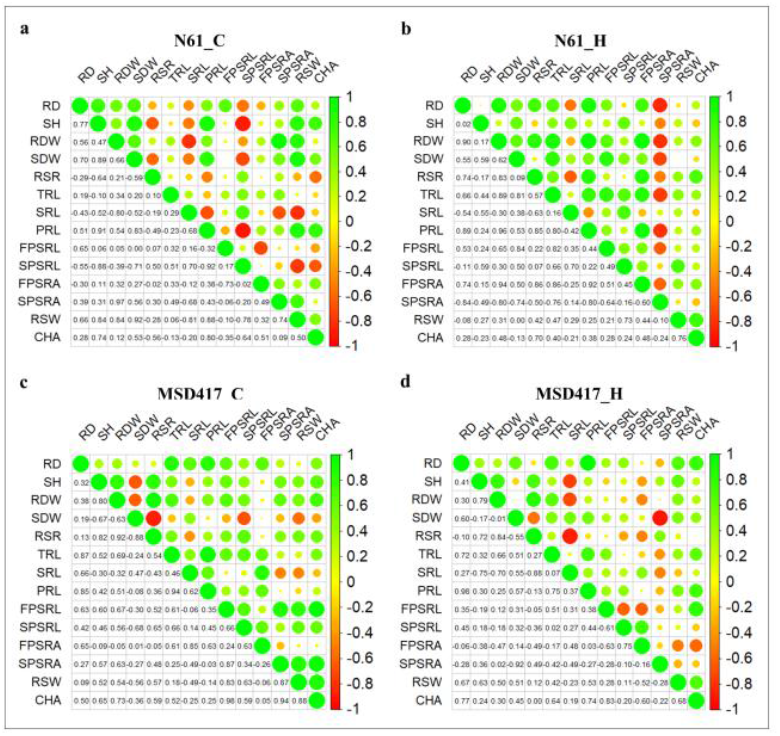
Positive correlation (in green) and negative correlation (in red) among different root and shoot traits. N61 in (a) control condition, and (b) high-temperature condition; and MSD417 in (c) control condition, and (d) high-temperature condition, n= 6. Abbreviations of the trait parameters are shown in Table 1

High temperature induced significant shifts in trait correlations across both genotypes (Fig.6). In N61, correlations involving SPSRA shifted from positive to strongly negative, while those involving SPSRL reversed from strongly negative to positive. A similar trend occurred in MSD417, where the correlation between RDW and SDW weakened significantly under high-temperature conditions, as the relationships of SRL with RDW, RSR, and SH, and that of SDW with SPSRA changed to strong negative correlations. Despite these shifts, certain relationships remained stable across all genotype-temperature condition combinations: CHA and RSW maintained a consistent positive correlation, whereas SRL consistently showed a negative correlation with RDW.

### Topology of the seminal root emergence

Analysis of root traits of the MSD417 described above suggested that this genotype has a larger angle of the outermost seminal roots, as indicated by the higher SPSRA value (Fig. 3i). To examine how this trait relates to seed morphogenesis during germination, the root emergence pattern from seeds was compared microscopically (Fig. 7). At seven days after germination, a massive clump of coleorhiza cells was observed at the base of the primary and other seminal roots in both genotypes (Figs. 7a, b). Depth of the initiation point of the outermost seminal root, estimated by calculating the distance between the intersection point of the perpendicular line from the center of the root initiation point and the root axis, and the boundary line between stem-root tissue, was statistically indistinguishable between N61 and MSD417 (Figs. 7c, d, e). However, N61 developed a coleorhiza tissue that extended vertically downward, unlike MSD417, forming a larger, lob-like structure (Figs. 7a, f). Consequently, MSD417 had a smaller height-to-width ratio of coleorhiza tissue compared to N61 (Fig. 7g), suggesting a suppression of coleorhiza tissue expansion in the downward direction in MSD417.

**Fig. 7.**
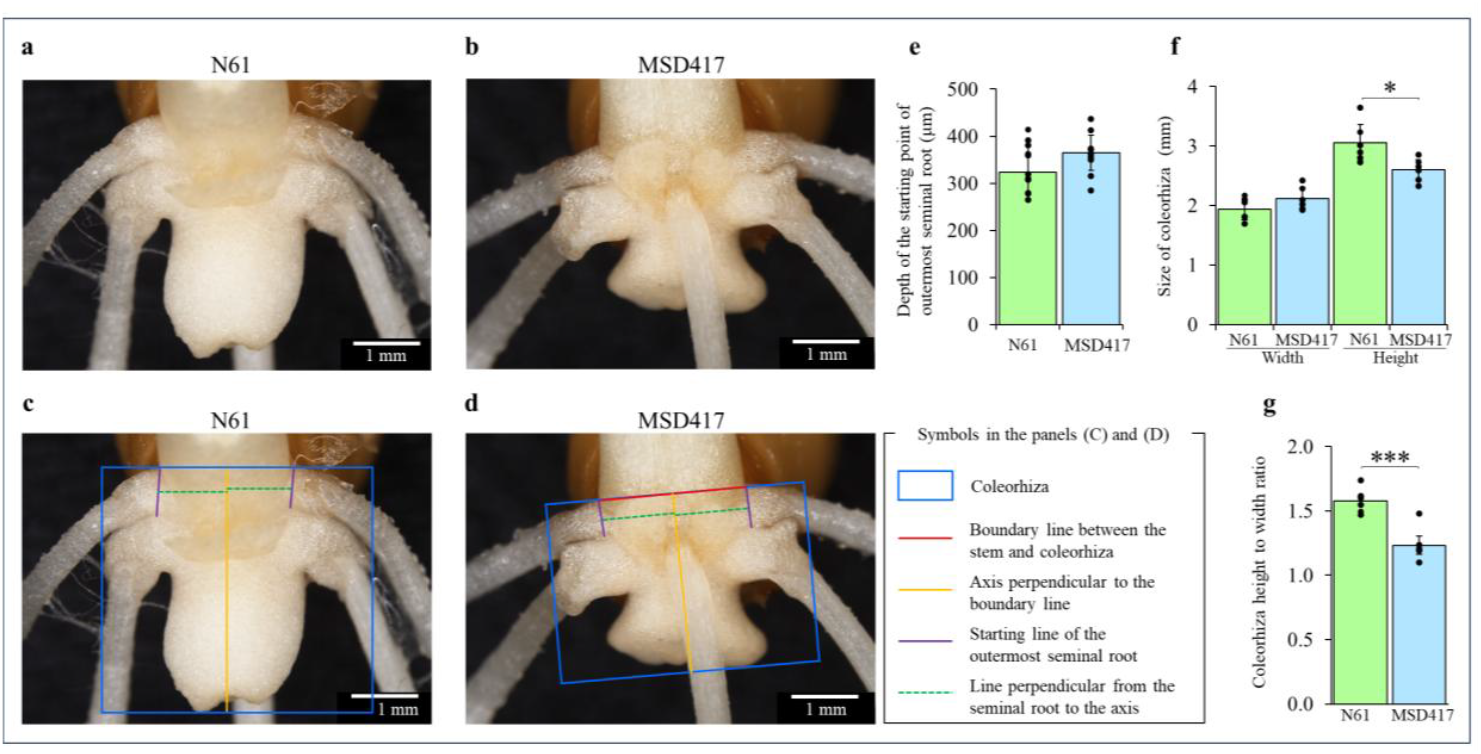
Microscopic representative images of N61 (a) and MSD417 (b) at 7 days after germination, grown in the control condition. Measuring technique of coleorhiza and depth of the starting point of the outermost seminal root of N61 (c), and MSD417 (d). Bar graph showing the depth of the starting point of the outermost seminal root (e), size of the coleorhiza (f), and coleorhiza height to width ratio of N61 and MSD417 at 7 days after germination (g). Data for each genotype were compared using Student’s t-test (p < 0.05, ***p < 0.001); n = 6

## Discussion

In this study, using a 2D-root phenotyping platform, we successfully characterized the *Aegilops tauschii*-derived MSD417 genotype as exhibiting a distinct root system architecture compared to its recurrent parent, N61, under controlled, high-temperature conditions. Under control conditions, MSD417 developed a larger root system, characterized by a wider angle of the outermost roots and a unique transverse expansion of the coleorhiza; the latter feature may suggest novel cell wall remodeling mechanisms, as discussed below. These results revealed that introgression from *Aegilops tauschii* into the MSD417 genotype resulted in distinct modifications in root system architecture, rather than changes in overall biomass accumulation or rooting depth. Despite the limitations imposed by high temperatures, the higher specific root length of MSD417 indicates an inherited advantage in resource acquisition.

Root system architecture (RSA) is one of the primary factors influencing crop productivity, significantly impacting global food supply through water use efficiency, soil nutrient uptake, and other agronomic traits **(Lynch 2007, 2022)**. Developing climate-resilient wheat genotypes that are better adapted to the challenges of climate change requires understanding how introgression alters RSA without compromising above-ground growth. Although wheat MSD population showed a wide variation for various agronomic traits, including grain yield, biomass, heading time, stress tolerance, resource use efficiency, and nutritional contents and composition **(Itam et al. 2021; Mohamed et al. 2022; Gorafi et al. 2025; Emam et al. 2025a, b)**their RSA characteristics have been poorly described.

Under the control condition, MSD417 exhibited a higher root-to-shoot ratio (RSR) and total root length (TRL) compared to N61, indicating that MSD417 allocated more resources to root growth (Fig.3c, f). Although rooting depth (RD) was not different between the two genotypes, the second pair of seminal root length (SPSRL) and root system width (RSW) were significantly larger in MSD417 than in N61, resulting in the larger convex hull area (CHA) in MSD417 (Figs. 3d, n, g, h). The larger RSW in MSD417 was also observed under high-temperature conditions (Fig. 3g), suggesting that MSD417 may possess a trait characterized by enhanced lateral root growth.

The larger angle of the outermost, second pair of seminal roots and the vigorous growth of these seminal roots during the juvenile stage suggested that the broader root system of MSD417 is at least partially attributable to the more lateral growth characteristics of the outermost seminal roots (Figs. 3j, n). The larger angle and enhanced growth of the second pair of seminal roots were also indicated by PCA biplot as prominent traits of MSD417 (Fig. 5). A diversity in the angle of the second pair of seminal roots has been reported in wheat cultivars grown in agar and soil systems **(Rich et al. 2020)**.

In many crops, the root angle significantly affects the efficiency of acquiring water and nutrients from different soil layers **(Kirschner et al. 2024)**. Deep root systems are considered superior at acquiring mobile nutrients such as water and nitrogen. On the other hand, shallow root systems formed by the horizontally expanded roots are thought to be advantageous for absorbing phosphorus, which has low mobility and tends to accumulate near the topsoil, as well as for adapting to waterlogging by obtaining oxygen for root respiration **(Mano and Omori 2007)**. The lateral root elongation is thought to involve the competing gravitropic and antigravitropic offset mechanisms **(Kirschner et al. 2024)**. A complex gene network has been proposed for gravity perception, signal transduction, and root morphological responses. For example, studies in molecular genetics and cellular biology showed that the actin-binding protein Rice Morphology Determinant (RMD) mediates the interaction between actin filaments and gravity-sensing organelles (statoliths) in rice roots, thereby regulating root growth angle in response to soil phosphate availability **(Huang et al. 2018)**. In other studies, rice mutants of DEEPER ROOTING 1 (*DRO1*) and the quantitative trait locus SOIL SURFACE ROOTING 1 (*qSOR1*), the homologues of LAZY genes responsible for root gravity sensing in Arabidopsis, exhibited reduced root gravitropism and resulted in a shallower root angle **(Uga et al. 2013; Kitomi et al. 2020; Oo et al. 2021)**. Therefore, the enhanced lateral root elongation observed in MSD417 in the present study suggests that this genotype may possess distinct genetic factors for root gravity sensing, signal transduction, or response mechanisms compared with its recurrent parent, N61.

In crop plants, a tradeoff between the allocation of root biomass near the soil surface and deep exploration has been discussed **(Maqbool et al. 2022)**. In the present study, it is noteworthy that MSD417 showed a larger RSW without reducing RD (Figs. 3g, d), resulting in a larger CHA than N61 (Fig. 3h). It has been suggested that CHA is an indicator for soil exploration to capture below-ground resources **(Bodner et al. 2019; Rangarajan and Lynch 2021)**. Enhanced biomass production and yield in MSD417 were reported in a previous study **(Matsunaga 2022)**. Further studies are needed to investigate the field performance of MSD417, including its ability to acquire water and nutrients from the underground, and its impact on yield.

This study showed that root growth of both MSD417 and N61 genotypes was markedly inhibited upon exposure to acute heat stress at a daytime temperature of 42 °C for four days (Fig. 3). No significant genotypic differences were found in most RSA parameters, such as root length and biomass, suggesting that the responses of both genotypes to high-temperature stress were largely similar, at least during the juvenile stage. Extremely high temperature stress has been known to inhibit root growth and significantly affect root system architecture **(Huang et al. 1991a, b; Pardales et al. 1999; Morales et al. 2003; Calleja-Cabrera et al. 2020)**. The cellular and molecular processes underlying heat-induced growth inhibition include decreased cellular membrane integrity in roots **(Iglesias-Acosta et al. 2010; Ionenko et al. 2010; Gupta et al. 2013)**, denaturation of cellular proteins and nucleic acids **(Howell 2013; Vu et al. 2019)**, increased production of reactive oxygen species and lipid peroxidation products **(Savicka and Škute 2010)**, decreased translocation of photoassimilates from the above ground tissues **(Heckathorn et al. 2013)**, leading to synergistically reduced rates of cell division and elongation in the root tissues **(Sattelmacher et al. 1990; Pardales et al. 1999; Joshi et al. 2016)**. The current study applied acute heat stress, which may have had determinant effects on RSA and obscured genotypic differences. Future studies aimed at examining the RSA response of MSD417 and the MSD population to high temperature should consider applying a gradual temperature increment and a less severe treatment to mimic field conditions.

Under high temperature conditions, two parameters, SRL and RSW, were found to be significantly different between MSD417 and N61. Moreover, MSD417 had a significantly wider SPSRA than N61. Among these parameters, SPSRA are considered to be inherited traits determined during the four days at controlled temperatures, rather than by their response to high temperature. Higher SRL in MSD417 under high-temperature conditions than that in N61 is intriguing: SRL is a commonly measured parameter, but its physiological significance remains to be fully resolved **(Freschet et al. 2021)**. High SRL may suggest thinner roots and a lower cost of root system construction, which requires less carbon and potentially leads to higher soil resource uptake efficiency. However, the cheaply constructed root system may have a shorter lifespan, thus it can be less efficient for long-term resource uptake **(Ryser 1996; Weemstra et al. 2020)**. The physiological significance of higher SRL in MSD417 should be examined in future studies.

The MSD417 exhibited a distinct coleorhiza morphological feature, which was larger than that of N61 at 7 days after germination (Fig. 7). Coleorhiza is a sheath-like, nonvascular organ that surrounds the radicle in grass embryos **(Holloway et al. 2021; Viana et al. 2022; Venglat et al. 2025)**. Coleorhiza is established before the end of seed development, and functions in the control of dormancy maintenance and germination through modulation of cell wall remodeling **(Holloway et al. 2021)**. Upon germination, the coleorhiza expands from the caryopsis into the environment, and then the radicle emerges into the soil by rupturing the coleorhiza. In *Avena fatua* (wild oat), coleorhiza growth occurs before the growth of the radicle, and is achieved by cell expansion mainly in the longitudinal direction **(Holloway et al. 2021)**. In this step, a localized expression of cell wall remodeling proteins, xyloglucan endotransglycosylases/ hydrolases, is implicated in this expansion. The tendency for longitudinal cell expansion was also observed in the present study in N61. In contrast, the transversally active expansion of coleorhiza in MSD417 suggests that cell wall remodeling may also be activated in the lateral side of the coleorhiza in this genotype. Since gene expression of cell wall remodeling proteins is regulated by plant hormones such as gibberellins and ethylene **(Ruduś et al. 2019; Calleja-Cabrera et al. 2020)**, the altered coleorhiza expansion in MSD417 may involve differences in the mechanisms of action of these plant hormones.

In the present study, we developed a versatile 2D root phenotyping system and characterized a unique wide root angle trait in the wheat MSD417 genotype. The multiple synthetic derivatives (MSD) population has over 400 representative genotypes **(Gorafi et al. 2025)**. Analysis of their root system architecture (RSA) in a high-throughput manner in the future may lead to the discovery of new genotypes exhibiting distinct root development traits for agricultural benefits. Moreover, although previous studies have indicated that the RSA traits at the seedling stage, including rooting angle, could be a useful proxy for those of mature root systems **(Maccaferri et al. 2016; Alahmad et al. 2019; Kang et al. 2024)**, verification is needed to determine whether the trait of enhanced lateral root development in MSD417 is also observed in the mature crop under field conditions. Overall, continued research on the genetic and environmental regulation of root development will contribute to a framework for improving the agricultural performance of wheat, thereby supporting the development of cultivars with improved productivity.

In this study, we successfully employed a simplified 2D-root phenotyping platform and characterized MSD417 as possessing distinct root traits compared with its recurrent parent, N61, under contrasting temperature regimes. Under control conditions, MSD417 developed a larger root system primarily as a consequence of an increased root system width, which was driven by a broader angle of the outermost pair of seminal roots, while maintaining comparable rooting depth to N61. This architectural configuration suggests that MSD417 may overcome the typical trade-off between shallow and deep soil foraging, a trait highly desirable for efficient nutrient and water capture. Furthermore, the unique transversal expansion of the coleorhiza in MSD417 may indicate potentially novel, genotype-specific mechanisms of cell wall remodeling that should be explored at the molecular level. Although high-temperature stress obscured the major morphological differences between genotypes, maintaining higher specific root length in MSD417 indicates an inherent capacity for efficient resource acquisition even under high-temperature stress. Nevertheless, to establish MSD417 as a valuable genotype for breeding programs, further validation of these traits in mature plants under field conditions is required. Ultimately, the experimental approaches presented here provide a framework for leveraging the genetic diversity of the MSD and similar populations to uncover variation in RSA.

## Supporting information

Supplementary-R-Script_Islam

## Supplementary Materials

The following supporting information is available: Document S1: R-scripts for the processing of data for root system architecture.

## Author Contributions

Conceptualization, S.M.M.I. and K.A.; methodology, S.M.M.I. and KA; software, S.M.M.I.; validation, S.M.M.I., I.S.A.T., and K.A.; formal analysis, S.M.M.I.; investigation, S.M.M.I.; resources, S.M.M.I.; data curation, S.M.M.I.; writing—original draft preparation, S.M.M.I.; writing—review and editing, I.S.A.T. and K.A.; visualization, S.M.M.I. and K.A.; supervision, K.A.; project administration, K.A.; funding acquisition, K.A. All authors have read and agreed to the published version of the manuscript.

## Data Availability Statement

The root images of wheat N61 and MSD417 are deposited in the Zenodo data repository under https://doi.org/10.5281/zenodo.18080159 and https://doi.org/10.5281/zenodo.18079748, respectively. The microscopic images of coleorhiza are deposited under https://doi.org/10.5281/zenodo.18091131. The other original contributions presented in the study are included in the article/supplementary material.

## Acknowledgments

We are grateful to Prof Hiroyuki Tanaka (Faculty of Agriculture, Tottori University, Tottori, Japan) and Dr Yasir Serag Alnor Gorafi (Graduate School of Agriculture, Kyoto University, Kyoto, Japan) for providing seeds of wheat cultivar Norin 61 and the MSD genotype MSD417, respectively. We thank the staff members of the Laboratory of Molecular and Cellular Biology, Faculty of Agriculture, Tottori University, for their technical support in the laboratory.

