## Supplementary-R-Script_Islam for "A practical phenotyping framework for root system architecture reveals enhanced root vigor in an *Aegilops tauschii*-derived wheat line"

**Supplementary1**
R-scripts for the processing of data for root system architecture

**Title:** A practical phenotyping framework for root system architecture reveals enhanced root vigor in an *Aegilops tauschii*-derived wheat line

Sultan Md Monwarul Islam[1], Izzat Sidahmed Ali Tahir[2,3], and Kinya Akashi [1,3,4 *]

1 United Graduate School of Agricultural Sciences, Tottori University, 4-101 Koyama-Cho-Minami, Tottori City, Tottori Prefecture, 680-0945, Japan;
2 Wheat Research Program, Agricultural Research Corporation, Wad Medani P.O. Box 126, Sudan;
3 International Platform for Dryland Research and Education, Tottori University, 1390 Hamasaka, Tottori City, Tottori Prefecture, 680-0001, Japan;
4 Faculty of Agriculture, Tottori University, 4-101 Koama-Chou-Minami, Tottori City, Tottori Prefecture, 680-0945, Japan;

**List of scripts:**
Script number 1. ANOVA and Tukeys HSD post hoc test for different root and shoot traits of N61 and MSD417 in contrasting temperature condition (Page: 01-02).
Script number 2. Student's t-test for different root and shoot traits of N61 and MSD417 in contrasting temperature condition (Page: 02-04).
Script number 3. ANOVA and Tukeys HSD post hoc test for different root traits of N61 and MSD417 from day 1 to day 8 in contrasting temperature conditions (Page: 04-11).
Script number 4. Principle component analysis (PCA) for different root and shoot traits of N61 and MSD417 in contrasting temperature conditions (Page: 11-15).
Script number 5. Correlation among different root and shoot traits of N61 and MSD417 in contrasting temperature condition (Page: 16-18).

Script number 1.
ANOVA and Tukeys HSD post hoc test for different root and shoot traits of N61 and MSD417 at day eight in contrasting temperature conditions.

```
# Install necessary packages
        install.packages("agricolae")
        install.packages("dplyr")
        install.packages("ggplot2")
        install.packages("broom")

# Load the packages
         library(agricolae)
         library(dplyr)
         library(ggplot2)
         library(broom)
         library(tidyverse)

# Clean up the system brain if any
         rm(list=ls())

# Select csv file by the following command
          setwd("D:/251006_R_analysis")
          FinalDayData <-read.csv("D:/251006_R_analysis/FinalDayData.csv")
```

```
Variables <- c("RD", "SH", "RDW", "SDW", "RSR","TRL", "SRL", "PRL", "FPSRL", "SPSRL", "RSW", 
"CHA", "FPSRA", "SPSRA")

ANOVA_FDD <- FinalDayData %>%
                        pivot_longer(
                        cols = all_of(Variables),
                        names_to = "Variable",
                        values_to = "Value") %>%
                        group_by(Variable) %>%
                        reframe(broom::tidy(aov(Value ~ Genotype))) %>%
                        filter(term == "Genotype") %>%
                        ungroup()

#Save the results to a CSV
                        write.csv(ANOVA_FDD, "ANOVA_FDD.csv", row.names = FALSE)


Tukeys HSD test
HSD_groups_FDD <-FinalDayData %>%
                          pivot_longer(
                          cols = all_of(Variables),
                          names_to = "Variable",
                          values_to = "Value") %>%
                          group_by(Variable) %>%
                          nest() %>%
                          mutate(model = map(data, ~ aov(Value ~ Genotype, data = .)),
                          hsd_result = map(model, ~ HSD.test(., trt = "Genotype", alpha = 0.05, console    = 
                          FALSE)),
                          groups_df = map(hsd_result, ~ as.data.frame(.$groups) %>%
                          rownames_to_column(var = "Genotype"))) %>%
                          select(Variable, groups_df) %>%
                          unnest(cols = groups_df) %>%
                          rename(Mean = Value)

#Save the results as CSV
write.csv(HSD_groups_FDD, "Tukeys_HSD_groups_FDD.csv", row.names = FALSE)
```

Script number 2.

<u>Student's t-test for different root and shoot traits of N61 and MSD417 in contrasting temperature conditions.</u>

```
#Install the necessary packages
        install.packages("agricolae")
        install.packages("dplyr")
        install.packages("ggplot2")
        install.packages("broom")
        install.packages("tidyverse")

# Load installed the packages
        library(agricolae)
        library(dplyr)
        library(ggplot2)
        library(broom)
        library(tidyverse)

# Memory cleaning and folder preparation
# (1) Clean up the system brain if any
        rm(list=ls())

# (2) Select csv file by the following command
        setwd("D:/251006_R_analysis")  # To select the desired location to save the file
```

```
        FinalDayData<-read.csv("D:/251006_R_analysis/FinalDayData.csv")

# t-test: N61_C vs. N61_H
# Create a subset data for N61 in two conditions
        subset_N61 <- subset(FinalDayData, Genotype %in% c("N61_C", "N61_H"))


Variables <- c("RD", "SH", "RDW", "SDW", "RSR","TRL", "SRL", "PRL", "FPSRL", "SPSRL",
            "RSW", "CHA", "FPSRA", "SPSRA")

# Run t-Test
        t_Test_N61 <- subset_N61 %>%
                        pivot_longer(
                        cols = all_of(Variables),
                        names_to = "Variable",
                        values_to = "Value") %>%
                        group_by(Variable) %>%
                        summarize(broom::tidy(t.test(Value ~ Genotype))) %>%
                        ungroup() %>%
                        rename(t_statistic = statistic,
                        p_value = p.value,
                        degrees_of_freedom = parameter,
                        conf_interval_low = conf.low,
                        conf_interval_high = conf.high,
                        mean_group1 = estimate1,
                        mean_group2 = estimate2)

# Save the results to a CSV
        write.csv(t_Test_N61, "t_Test_N61_results.csv", row.names = FALSE)


# t-test: MSD417_C vs. MSD417_H
        subset_MSD417 <- subset(FinalDayData, Genotype %in% c("MSD417_C", "MSD417_H"))

# Run t-Test
        t_Test_MSD417 <- subset_MSD417 %>%
                          pivot_longer(
                          cols = all_of(Variables),
                          names_to = "Variable",
                          values_to = "Value") %>%
                          group_by(Variable) %>%
                          summarize(broom::tidy(t.test(Value ~ Genotype))) %>%
                          ungroup() %>%
                          rename(t_statistic = statistic,
                          p_value = p.value,
                          degrees_of_freedom = parameter,
                          conf_interval_low = conf.low,
                           conf_interval_high = conf.high,
                           mean_group1 = estimate1,
                           mean_group2 = estimate2)

# Save the results as CSV
        write.csv(t_Test_MSD417, "t_Test_MSD417_results.csv", row.names = FALSE)
```

Script number 3.
<u>ANOVA and Tukeys HSD post hoc test for different root traits of N61 and MSD417 from day 1 to day 4 in contrasting temperature conditions.</u>

```
# Necessary packages
        install.packages("agricolae")
        install.packages("dplyr")
        install.packages("ggplot2")
        install.packages("broom")

# Load the packages
        library(agricolae)
        library(dplyr)
        library(ggplot2)
        library(broom)
        library(tidyverse)

# Memory cleaning and folder preparation
# (1) Clean up the system brain if any
         rm(list=ls())

# (2) Select csv file by the following command
        setwd("D:/251006_R_analysis")
        DailyData <-read.csv("D:/251006_R_analysis/DailyData_all.csv")

#ANOVA and Tukeys HSD: Day1
# Create a sub-set data for day 1
        Data_day1 <- subset(DailyData, Days %in% c("Day1"))

# ANOVA preparation
        Variables <- c("PRL", "FPSRL", "SPSRL", "TRL", "RD", "RSW", "CHA")

        ANOVA_Day1 <- Data_day1 %>%
                        pivot_longer(
                        cols = all_of(Variables),
                        names_to = "Variable",
                        values_to = "Value") %>%
                        filter(Variable != "SPSRL") %>%
                        group_by(Variable) %>%
                        reframe(broom::tidy(aov(Value ~ Genotype))) %>%
                        filter(term == "Genotype") %>%
        ungroup()

# Save the results to a CSV
        write.csv(ANOVA_Day1, "ANOVA_Day1.csv", row.names = FALSE)

# Tukeys HSD test:  Day1
        HSD_groups_Day1 <-Data_day1 %>%
                pivot_longer(
                cols = all_of(Variables),
                names_to = "Variable",
                values_to = "Value") %>%
                group_by(Variable) %>%
                filter(Variable != "SPSRL") %>%
                nest() %>%
                mutate(model = map(data, ~ aov(Value ~ Genotype, data = .)),
                hsd_result = map(model, ~ HSD.test(., trt = "Genotype",
                alpha = 0.05, console = FALSE)),
                groups_df = map(hsd_result, ~ as.data.frame(.$groups) %>%
                rownames_to_column(var = "Genotype"))) %>%
                select(Variable, groups_df) %>%
                unnest(cols = groups_df) %>%
                rename(Mean = Value)
```

```
# Save the results as CSV
        write.csv(HSD_groups_Day1, "Tukeys_HSD_groups_Day1.csv", row.names = FALSE)


#ANOVA and Tukeys HSD: Day2
# Create a sub-set data for day 2
        Data_day2 <- subset(DailyData, Days %in% c("Day2"))

# ANOVA preparation
        ANOVA_Day2 <- Data_day2 %>%
                        pivot_longer(
                        cols = all_of(Variables),
                        names_to = "Variable",
                        values_to = "Value") %>%
                        group_by(Variable) %>%
                        reframe(broom::tidy(aov(Value ~ Genotype))) %>%
                        filter(term == "Genotype") %>%
                        ungroup()

# Save the results to a CSV
        write.csv(ANOVA_Day2, "ANOVA_Day2.csv", row.names = FALSE)

# Tukeys HSD test:  Day2
        HSD_groups_Day2 <-Data_day2 %>%
                pivot_longer(
                cols = all_of(Variables),
                names_to = "Variable",
                values_to = "Value") %>%
                group_by(Variable) %>%
                nest() %>%
                mutate(model = map(data, ~ aov(Value ~ Genotype, data = .)),
                hsd_result = map(model, ~ HSD.test(., trt = "Genotype",
                alpha = 0.05, console = FALSE)),
                groups_df = map(hsd_result, ~ as.data.frame(.$groups) %>%
                rownames_to_column(var = "Genotype"))) %>%
                select(Variable, groups_df) %>%
                unnest(cols = groups_df) %>%
                rename(Mean = Value)

# Save the results as CSV
        write.csv(HSD_groups_Day2, "Tukeys_HSD_groups_Day2.csv", row.names = FALSE)


#ANOVA and Tukeys HSD: Day3
# Create a sub-set data for day 3
        Data_day3 <- subset(DailyData, Days %in% c("Day3"))

# ANOVA preparation
        ANOVA_Day3 <- Data_day3 %>%
                        pivot_longer(
                        cols = all_of(Variables),
                        names_to = "Variable",
                        values_to = "Value") %>%
                        group_by(Variable) %>%
                        reframe(broom::tidy(aov(Value ~ Genotype))) %>%
                        filter(term == "Genotype") %>%
                        ungroup()

# Save the results to a CSV
        write.csv(ANOVA_Day3, "ANOVA_Day3.csv", row.names = FALSE)
```

```
# Tukeys HSD test:  Day3
          HSD_groups_Day3 <-Data_day3 %>%
                    pivot_longer(
                    cols = all_of(Variables),
                    names_to = "Variable",
                    values_to = "Value") %>%
                    group_by(Variable) %>%
                    nest() %>%
                    mutate(model = map(data, ~ aov(Value ~ Genotype, data = .)),
                    hsd_result = map(model, ~ HSD.test(., trt = "Genotype",
                    alpha = 0.05, console = FALSE)),
                    groups_df = map(hsd_result, ~ as.data.frame(.$groups) %>%
                    rownames_to_column(var = "Genotype"))) %>%
                    select(Variable, groups_df) %>%
                    unnest(cols = groups_df) %>%
                    rename(Mean = Value)

# Save the results as CSV
          write.csv(HSD_groups_Day3, "Tukeys_HSD_groups_Day3.csv", row.names = FALSE)


#ANOVA and Tukeys HSD: Day4
# Create a sub-set data for day 4
          Data_day4 <- subset(DailyData, Days %in% c("Day4"))

# ANOVA preparation
          ANOVA_Day4 <- Data_day4 %>%
                              pivot_longer(
                              cols = all_of(Variables),
                              names_to = "Variable",
                              values_to = "Value") %>%
                              group_by(Variable) %>%
                              reframe(broom::tidy(aov(Value ~ Genotype))) %>%
                              filter(term == "Genotype") %>%
                              ungroup()

# Save the results to a CSV
          write.csv(ANOVA_Day4, "ANOVA_Day4.csv", row.names = FALSE)

# Tukeys HSD test:  Day4
          HSD_groups_Day4 <-Data_day4 %>%
                    pivot_longer(
                    cols = all_of(Variables),
                    names_to = "Variable",
                    values_to = "Value") %>%
                    group_by(Variable) %>%
                    nest() %>%
                    mutate(model = map(data, ~ aov(Value ~ Genotype, data = .)),
                    hsd_result = map(model, ~ HSD.test(., trt = "Genotype",
                    alpha = 0.05, console = FALSE)),
                    groups_df = map(hsd_result, ~ as.data.frame(.$groups) %>%
                    rownames_to_column(var = "Genotype"))) %>%
                    select(Variable, groups_df) %>%
                    unnest(cols = groups_df) %>%
                    rename(Mean = Value)

# Save the results as CSV
          write.csv(HSD_groups_Day4, "Tukeys_HSD_groups_Day4.csv", row.names = FALSE)
```

#ANOVA and Tukeys HSD: Day5
### Create a sub-set data for day 5

```
Data_day5 <- subset(DailyData, Days %in% c("Day5"))
```

### ANOVA preparation

```
ANOVA_Day5 <- Data_day5 %>%
        pivot_longer(
        cols = all_of(Variables),
        names_to = "Variable",
        values_to = "Value") %>%
        group_by(Variable) %>%
        reframe(broom::tidy(aov(Value ~ Genotype))) %>%
        filter(term == "Genotype") %>%
        ungroup()
```

### Save the results to a CSV

```
write.csv(ANOVA_Day5, "ANOVA_Day5.csv", row.names = FALSE)
```

### Tukeys HSD test:  Day5

```
HSD_groups_Day5 <-Data_day5 %>%
    pivot_longer(
    cols = all_of(Variables),
    names_to = "Variable",
    values_to = "Value") %>%
    group_by(Variable) %>%
    nest() %>%
    mutate(model = map(data, ~ aov(Value ~ Genotype, data = .)),
    hsd_result = map(model, ~ HSD.test(., trt = "Genotype",
    alpha = 0.05, console = FALSE)),
    groups_df = map(hsd_result, ~ as.data.frame(.$groups) %>%
    rownames_to_column(var = "Genotype"))) %>%
    select(Variable, groups_df) %>%
    unnest(cols = groups_df) %>%
    rename(Mean = Value)
```

### Save the results as CSV

```
write.csv(HSD_groups_Day5, "Tukeys_HSD_groups_Day5.csv", row.names = FALSE)
```

#ANOVA and Tukeys HSD: Day6
### Create a sub-set data for day 6

```
Data_day6 <- subset(DailyData, Days %in% c("Day6"))
```

### ANOVA preparation

```
ANOVA_Day6 <- Data_day6 %>%
        pivot_longer(
        cols = all_of(Variables),
        names_to = "Variable",
        values_to = "Value") %>%
        group_by(Variable) %>%
        reframe(broom::tidy(aov(Value ~ Genotype))) %>%
        filter(term == "Genotype") %>%
        ungroup()
```

### Save the results to a CSV

```
write.csv(ANOVA_Day6, "ANOVA_Day6.csv", row.names = FALSE)
```

### Tukeys HSD test:  Day6

```
HSD_groups_Day6 <-Data_day6 %>%
```

```
                  pivot_longer(
                  cols = all_of(Variables),
                  names_to = "Variable",
                  values_to = "Value") %>%
                  group_by(Variable) %>%
                  nest() %>%
                  mutate(model = map(data, ~ aov(Value ~ Genotype, data = .)),
                  hsd_result = map(model, ~ HSD.test(., trt = "Genotype",
                  alpha = 0.05, console = FALSE)),
                  groups_df = map(hsd_result, ~ as.data.frame(.$groups) %>%
                  rownames_to_column(var = "Genotype"))) %>%
                  select(Variable, groups_df) %>%
                  unnest(cols = groups_df) %>%
                  rename(Mean = Value)

# Save the results as CSV
      write.csv(HSD_groups_Day6, "Tukeys_HSD_groups_Day6.csv", row.names = FALSE)


#ANOVA and Tukeys HSD: Day7
# Create a sub-set data for day 7
      Data_day7 <- subset(DailyData, Days %in% c("Day7"))

# ANOVA preparation
      ANOVA_Day7 <- Data_day7 %>%
                        pivot_longer(
                        cols = all_of(Variables),
                        names_to = "Variable",
                        values_to = "Value") %>%
                        group_by(Variable) %>%
                        reframe(broom::tidy(aov(Value ~ Genotype))) %>%
                        filter(term == "Genotype") %>%
                        ungroup()

# Save the results to a CSV
      write.csv(ANOVA_Day7, "ANOVA_Day7.csv", row.names = FALSE)

# Tukeys HSD test:  Day7
      HSD_groups_Day7 <-Data_day7 %>%
                  pivot_longer(
                  cols = all_of(Variables),
                  names_to = "Variable",
                  values_to = "Value") %>%
                  group_by(Variable) %>%
                  nest() %>%
                  mutate(model = map(data, ~ aov(Value ~ Genotype, data = .)),
                  hsd_result = map(model, ~ HSD.test(., trt = "Genotype",
                  alpha = 0.05, console = FALSE)),
                  groups_df = map(hsd_result, ~ as.data.frame(.$groups) %>%
                  rownames_to_column(var = "Genotype"))) %>%
                  select(Variable, groups_df) %>%
                  unnest(cols = groups_df) %>%
                  rename(Mean = Value)

# Save the results as CSV
      write.csv(HSD_groups_Day7, "Tukeys_HSD_groups_Day7.csv", row.names = FALSE)

#ANOVA and Tukeys HSD: Day8
# Create a sub-set data for day 8
      Data_day8 <- subset(DailyData, Days %in% c("Day8"))
```

```
# ANOVA preparation
        ANOVA_Day8 <- Data_day8%>%
                    pivot_longer(
                    cols = all_of(Variables),
                    names_to = "Variable",
                    values_to = "Value") %>%
                    group_by(Variable) %>%
                    reframe(broom::tidy(aov(Value ~ Genotype))) %>%
                    filter(term == "Genotype") %>%
                    ungroup()

# Save the results to a CSV
        write.csv(ANOVA_Day8, "ANOVA_Day8.csv", row.names = FALSE)

# Tukeys HSD test:  Day8
        HSD_groups_Day8 <-Data_day8 %>%
                pivot_longer(
                cols = all_of(Variables),
                names_to = "Variable",
                values_to = "Value") %>%
                group_by(Variable) %>%
                nest() %>%
                mutate(model = map(data, ~ aov(Value ~ Genotype, data = .)),
                hsd_result = map(model, ~ HSD.test(., trt = "Genotype",
                alpha = 0.05, console = FALSE)),
                groups_df = map(hsd_result, ~ as.data.frame(.$groups) %>%
                rownames_to_column(var = "Genotype"))) %>%
                select(Variable, groups_df) %>%
                unnest(cols = groups_df) %>%
                rename(Mean = Value)

# Save the results as CSV
        write.csv(HSD_groups_Day8, "Tukeys_HSD_groups_Day8.csv", row.names = FALSE)
```

Script number 4.

Principle component analysis (PCA) for different root and shoot traits of N61 and MSD417 in contrasting tempe rature conditions.

```
# Necessary packages
install.packages("conflicted")
install.packages("dplyr")
install.packages("ggplot2")
install.packages("readr")
install.packages("psych")
install.packages("ellipse")
install.packages("ggfortify")
install.packages("ggrepel")
install.packages("factoextra")
install.packages("RColorBrewer")
install.packages("FactoMineR")

# Load necessary packages
library(dplyr)
library(ggplot2)
library(readr)
library(psych)
library(ellipse)
```

```
library(ggfortify)
library(ggrepel)
library(factoextra)
library(RColorBrewer)
library(FactoMineR)

# Memory cleaning and folder preparation
Clean up the system memory
rm(list=ls())

# Select desired location to save the files
setwd("D:/251006_R_analysis")

# Give a new name and show the csv file path
FinalDayData<-read.csv("D:/251006_R_analysis/FinalDayData.csv")

# Separate numeric columns from the data set
PCA_numeric <- FinalDayData %>% select(4:17)

# Separate category info or Genotype column
PCA_id <- dplyr::select(FinalDayData, Genotype)

# Run PCA
PCA_result <- prcomp(PCA_numeric, center = TRUE, scale. = TRUE)

# Save PCA result data as specific data format as csv
PCA_result_download <-  as.data.frame(PCA_result$rotation)
pca_result_data <- paste0(format(Sys.Date(), "%y%m%d"),"_PCA_result_N61_MSD417", ".csv")
pca_result_data <- file.path("D:/251006_R_analysis", pca_result_data)
write.csv(PCA_result_download,pca_result_data, row.names=FALSE)

# Preparation of PCA score
PCA_scoreonly <-  as.data.frame(PCA_result$x)

# Combining the PCA_scoreonly with the ID, the genotype column
PCA_score <- cbind(PCA_id , PCA_scoreonly)

# Export the PCA score data as csv
PCA_score_data <- paste0(format(Sys.Date(), "%y%m%d"),"_PCA_score_N61_MSD417", ".csv")
PCA_score_data <- file.path("D:/251006_R_analysis", PCA_score_data)
write.csv(PCA_score, PCA_score_data, row.names=FALSE)

# Calculate and export stdev data as csv
PCA_stdev_only <- as.data.frame(PCA_result$sdev)
PCA_stdev <- paste0(format(Sys.Date(), "%y%m%d"),"_PCA_stdev_N61_MSD417", ".csv")
PCA_stdev <- file.path("D:/251006_R_analysis", PCA_stdev)
write.csv(PCA_stdev_only, PCA_stdev, row.names=FALSE)

# Calculate the loadings, and export as csv file
PCA_loadings_data <- sweep(PCA_result$rotation, MARGIN=2, PCA_result$sdev, FUN="*")
PCA_loading <- paste0(format(Sys.Date(), "%y%m%d"),"_PCA_loading_N61_MSD417", ".csv")
PCA_loading <- file.path("D:/251006_R_analysis", PCA_loading)
write.csv(PCA_loadings_data,PCA_loading, row.names=FALSE)

# Variance explained by each PC
PCA_varience <- PCA_result$sdev^2
PCA_varience_eachpc <- paste0(format(Sys.Date(), "%y%m%d"),"_PCA_varienc_N61_MSD417", ".csv")
PCA_varience_eachpc<- file.path("D:/251006_R_analysis", PCA_varience_eachpc)
write.csv(PCA_varience,PCA_varience_eachpc, row.names=FALSE)

# Percent varience explained by each PC
```

```
PCA_percent_varience <- round(100 * PCA_varience / sum(PCA_varience), 1)
PCA_percent_varience_data <- paste0(format(Sys.Date(), "%y%m%d"),"_PCA_Percent_varienc_N61_MSD41
7", ".csv")
PCA_percent_varience_data<- file.path("D:/251006_R_analysis", PCA_percent_varience_data)
write.csv(PCA_percent_varience, PCA_percent_varience_data, row.names=FALSE)

# Select the highest contributing variables:
Get loadings (contributions of variables to each PC)
loadings_T10 <- PCA_result$rotation #same as raw PCA_result, with variable name
write.csv(loadings_T10, file = "PCA_result_with_variable_name_N61_MSD417.csv")

# Compute contribution of each variable to PC1 and PC2
PCA_loading_strength_T10 <- rowSums(abs(loadings_T10[, 1:2]))
write.csv(PCA_loading_strength_T10, file = "PCA_loadings_strength_T10.csv")

# Separate top 10 contributing variables
top10_variables <- names(sort(PCA_loading_strength_T10, decreasing = TRUE))[1:10]
write.csv(top10_variables, file = "Top10_PCA_contributing_variables.csv")

# Separate the data for top 10 PCA contributing variables from the main data set
PCA_data_T10 <- PCA_numeric[, top10_variables]
Rerun PCA from the separated data of top 10 PCA contributing variables
PCA_result_T10 <- prcomp(PCA_data_T10, center = TRUE, scale. = TRUE)

# Save PCA result data as specific data format as csv
PCA_result_download_T10 <-  as.data.frame(PCA_result_T10$rotation)
pca_result_data_T10 <- paste0(format(Sys.Date(), "%y%m%d"),"_PCA_result_T10_N61_MSD417", ".csv")

pca_result_data_T10 <- file.path("D:/251006_R_analysis", pca_result_data_T10)
write.csv(PCA_result_download_T10,pca_result_data_T10, row.names=FALSE)

# Preparation of PCA score
PCA_scoreonly_T10 <-  as.data.frame(PCA_result_T10$x)

# Combining the PCA_scoreonly with the ID, the genotype column
PCA_score_T10 <- cbind(PCA_id , PCA_scoreonly_T10)

# Export the PCA score data as csv
PCA_score_data_T10 <- paste0(format(Sys.Date(), "%y%m%d"),"_PCA_score_T10_N61_MSD417", ".csv")
PCA_score_data_T10 <- file.path("D:/251006_R_analysis", PCA_score_data_T10)
write.csv(PCA_score_T10, PCA_score_data_T10, row.names=FALSE)

# Calculate and export stdev data as csv
PCA_stdev_only_T10 <- as.data.frame(PCA_result_T10$sdev)
PCA_stdev_T10 <- paste0(format(Sys.Date(), "%y%m%d"),"_PCA_stdev_T10_N61_MSD417", ".csv")
PCA_stdev_T10 <- file.path("D:/251006_R_analysis", PCA_stdev_T10)
write.csv(PCA_stdev_only_T10, PCA_stdev_T10, row.names=FALSE)

# Calculate the loadings, and export as csv file
PCA_loadings_data_T10 <- sweep(PCA_result_T10$rotation, MARGIN=2, PCA_result_T10$sdev, FUN="*")
PCA_loading_T10 <- paste0(format(Sys.Date(), "%y%m%d"),"_PCA_loading_T10_N61_MSD417", ".csv")
PCA_loading <- file.path("D:/251006_R_analysis", PCA_loading)
write.csv(PCA_loadings_data_T10,PCA_loading_T10, row.names=FALSE)

# Variance explained by each PC
PCA_varience_T10 <- PCA_result_T10$sdev^2
PCA_varience_eachpc_T10 <- paste0(format(Sys.Date(), "%y%m%d"),"_PCA_varienc_T10_N61_MSD417",
".csv")
PCA_varience_eachpc_T10<- file.path("D:/251006_R_analysis", PCA_varience_eachpc_T10)
write.csv(PCA_varience_T10,PCA_varience_eachpc_T10, row.names=FALSE)
```

```
# Percent varience explained by each PC: percent of PCA varience
PCA_percent_varience_T10 <- round(100 * PCA_varience_T10 / sum(PCA_varience_T10), 1)
PCA_percent_varience_data_T10 <- paste0(format(Sys.Date(), "%y%m%d"),"_PCA_Percent_varienc_T10_N61_MSD417", ".csv")
PCA_percent_varience_data_T10<- file.path("D:/251006_R_analysis", PCA_percent_varience_data_T10)
write.csv(PCA_percent_varience_T10, PCA_percent_varience_data_T10, row.names=FALSE)

# Preparation of PCA plot based on PC1 and PC2
Separate the PCA score for PC1 and PC2
PC1_score_T10 <-  PCA_result_T10$x[,1]
PC2_score_T10 <-  PCA_result_T10$x[,2]

# Create PCA score data frame of separated PC score
PCA_scores_df_T10 <- as.data.frame(PCA_result_T10$x[, 1:2])
PCA_scores_df_T10$Genotype <-FinalDayData$Genotype

# Add variables contribution arrows
Separate PCA result for PC1 and PC2
PCA_loadings_df_T10 <- as.data.frame(PCA_result_T10$rotation[, 1:2])
PCA_loadings_df_T10$Variable <- rownames(PCA_loadings_df_T10)

# Add scale arrows to match the PCA score space
arrow_scale <- 8
PCA_loadings_df_T10$PC1 <- PCA_loadings_df_T10$PC1 * arrow_scale
PCA_loadings_df_T10$PC2 <- PCA_loadings_df_T10$PC2 * arrow_scale

PCA_plot_T10 <-ggplot(PCA_scores_df_T10, aes(x = PC1, y = PC2, color = Genotype)) +
        geom_point(size = 2.5) + stat_ellipse(aes(group = Genotype), linetype = "dashed", linewidth = 1.0) + scale_color_manual(values = c("N61_C" = "#28A800", "N61_H" = "#EAB200" ,"MSD417_C" = "#0066FF", "MSD417_H" = "#FF00FF")) + geom_segment(data = PCA_loadings_df_T10, aes(x = 0, y = 0, xend = PC1, yend = PC2), arrow = arrow(length = unit(0.2, "cm")), color = "black") + geom_text_repel(data = PCA_loadings_df_T10, aes(x = PC1, y = PC2, label = Variable), color = "black",  size = 5, box.padding = unit(0.1, "lines"), min.segment.length = unit(0, "lines"), max.overlaps = Inf,           segment.alpha = 0)+ theme_classic() + labs(title = "", x = paste0("PC1 (", PCA_percent_varience_T10[1], "%)"), y = paste0("PC2 (", PCA_percent_varience_T10[2], "%)")) + guides(color = guide_legend(override.aes = list(linetype = 0))) +theme_classic() + theme(axis.line = element_line(color = "black", linewidth = 0.5), axis.text = element_text(size = 16), axis.title = element_text(size = 18), legend.text = element_text(size = 14), legend.position = "inside", legend.position.inside = c(1.0, 1.0), legend.justification = c("left", "top"), legend.direction = "horizontal", legend.title = element_blank(), legend.background = element_blank(), panel.grid.major = element_line(color = "grey92", linewidth = 0.1, linetype = "dashed"), panel.grid.minor = element_line(color = "grey92", linewidth = 0.1, linetype = "dashed")) # Make legend background transparent

Display the plot
print(PCA_plot_T10)
```

Script number 5.

<u>Correlation among different root and shoot traits of N61 and MSD417 in contrasting temperature condition.</u>

*# Necessary packages*

```
install.packages("conflicted")
install.packages("dplyr")
install.packages("ggplot2")
install.packages("corrplot")
install.packages("ggcorrplot")
```

*# Load necessary packages*

```
library(conflicted)
```

```
library(dplyr)
library(ggplot2)
library(corrplot)
library(ggcorrplot)

# Memory cleaning and folder preparation
# Clean up the system memory
rm(list=ls())

# Select desired location to save the files
setwd("D:/251006_R_analysis")

# Give a new name and show the csv file path
FinalDayCorrelation<-read.csv("D:/251006_R_analysis/FinalDayData.csv")

# Exclude numeric columns
numeric_vars <- c("RD", "SH", "RDW", "SDW", "RSR", "TRL", "SRL", "PRL", "FPSRL", "SPSRL", "FPSRA
", "SPSRA", "RSW", "CHA")

# Separate data by treatment
N61_C <- FinalDayCorrelation[FinalDayCorrelation$Genotype == "N61_C", numeric_vars]
MSD417_C <- FinalDayCorrelation[FinalDayCorrelation$Genotype == "MSD417_C", numeric_vars]
N61_H <- FinalDayCorrelation[FinalDayCorrelation$Genotype == "N61_H", numeric_vars]
MSD417_H <- FinalDayCorrelation[FinalDayCorrelation$Genotype == "MSD417_H", numeric_vars]

# Calculate correlation matrices
cor_matrix_N61_C <- cor(N61_C, use = "pairwise.complete.obs")
cor_matrix_MSD417_C <- cor(MSD417_C, use = "pairwise.complete.obs")
cor_matrix_N61_H <- cor(N61_H, use = "pairwise.complete.obs")
cor_matrix_MSD417_H <- cor(MSD417_H, use = "pairwise.complete.obs")

# Set up the plotting layout to have two plots in one row
par(mfrow = c(2, 2), mar = c(1, 1, 3, 1))
Reset plotting layout to default
par(mfrow = c(1, 1), mar = c(5, 4, 4, 2) + 0.1)
Preaparation of correlation plot
my_colors <- colorRampPalette(c("red", "yellow", "green"))(200)


# Correlation plot: N61 in control condition (N61_C)

# Step 1: Plot the upper triangle with circles
corrplot(cor_matrix_N61_C, method = "circle", type = "upper", order = "original", col = my_colors, diag = TR
UE, tl.pos = "t", tl.col = "black", tl.srt = 45, tl.offset = 1.0, main = "Correlations for N61 in control condition",
mar = c(2,0,2,0), cl.align.text = "l", tl.cex = 1.5, cl.cex = 1.8)

# Step 2: Add the lower triangle with numeric coefficients
corrplot(cor_matrix_N61_C, add = TRUE, type = "lower", method = "number", order = "original",diag = FALS
E, col = "black", number.cex = 0.9, number.font = 1, srt = 45, tl.pos = "n", cl.pos = "n")

# Correlation plot: N61 in heat condition (N61_H)
# Step 1: Plot the upper triangle with circles
corrplot(cor_matrix_N61_H, method = "circle", type = "upper", order = "original", col = my_colors, diag = TR
UE, tl.pos = "t", tl.col = "black", tl.srt = 45, tl.offset = 1.0, main = "Correlations for N61 in heat condition", mar
= c(2,0,2,0), cl.align.text = "l", tl.cex = 1.5, cl.cex = 1.8)

# Step 2: Add the lower triangle with numeric coefficients
corrplot(cor_matrix_N61_H, add = TRUE, type = "lower", method = "number", order = "original", diag = FALS
E, col = "black", number.cex = 0.9, number.font = 1, srt = 45, tl.pos = "n", cl.pos = "n")

# Correlation plot: MSD417 in control condition (MSD417_C)
```

```
# Step 1: Plot the upper triangle with circles
corrplot(cor_matrix_MSD417_C, method = "circle", type = "upper", order = "original", col = my_colors, diag =
TRUE, tl.pos = "t", tl.col = "black", tl.srt = 45, tl.offset = 1.0, main = "Correlations for MSD417 in control cond
ition", mar = c(2,0,2,0), cl.align.text = "l", tl.cex = 1.5, cl.cex = 1.8)

# Step 2: Add the lower triangle with numeric coefficients
corrplot(cor_matrix_MSD417_C, add = TRUE, type = "lower", method = "number", order = "original", diag = F
ALSE, col = "black", number.cex = 0.9, number.font = 1, srt = 45, tl.pos = "n", cl.pos = "n")

# Correlation plot: MSD417 in heat condition (MSD417_H)
# Step 1: Plot the upper triangle with circles
corrplot(cor_matrix_MSD417_H, method = "circle", type = "upper", order = "original", col = my_colors, diag =
 TRUE, tl.pos = "t", tl.col = "black", tl.srt = 45, tl.offset = 1.0, main = "Correlations for MSD417 in heat conditi
on", mar = c(2,0,2,0), cl.align.text = "l", tl.cex = 1.5, cl.cex = 1.8)

# Step 2: Add the lower triangle with numeric coefficients
corrplot(cor_matrix_MSD417_H, add = TRUE, type = "lower", method = "number", order = "original", diag =
FALSE, col = "black", number.cex = 0.9, number.font = 1, srt = 45, tl.pos = "n", cl.pos = "n")
```
